# Genomic prediction for general combining ability in hybrid canola (*Brassica napa* L.)

**DOI:** 10.1101/2025.11.03.686416

**Authors:** Biructawit B. Tessema, C.P. Andrahennadi, Kathleen Poff, Kening Yao, Zoe Ehlert, Tom Kubik, Travis Anton, Kate Light, Wayne Burton, Kelly R Robbins

## Abstract

Hybrid breeding is a method of selecting parental lines and determining crosses that are likely to yield the best hybrids. Genomic prediction (GP), a tool that uses genome wide markers, can be employed to predict the performance of untested hybrids based on general combining ability (GCA) of their parents. We investigated the potential of GP for GCA prediction in a commercial canola breeding program. We used female tester data, where many female lines are crossed with few male lines, to predict economically important traits in canola. Multi-year and location data for grain yield, oil, protein, days to flowering, days to maturity, total glucosinate and saturated fat were available for prediction. Three different cross-validation strategies were implemented to determine the predictive ability (PA) of each trait. In the first cross-validation scheme (CV1), a prediction model was validated using five-fold cross validation strategy. In the second cross-validation scheme (CV2), an unseen year was predicted using the previous years’ data as a training set. In the third cross-validation scheme (CV3), Inbred pe se performance was added to the training set to exploit the covariance between traits of inbred and hybrid trials. The highest PA was observed for CV1 while the lowest PA was seen for CV2. In CV1, PA ranged from 0.34 to 0.62. The highest PA was observed for protein (0.62) while the lowest PA was observed for days to maturity (0.34). For CV2, PA ranged from 0.16 to 0.46, while for CV3 PA ranged from 0.27 to 0.71. The highest PA for CV2 (0.46) and CV3 (0.71) was observed for total glucosinate while the lowest PA (0.16 CV2 and 0.27 -CV3) was for days to maturity. The current study demonstrates the potential of using marker information to select parents with the highest GCA to create the best hybrid combinations.

## Introduction

Canola (*Brassica napus* L.) is an economically important oil crop (Qian *et al*.), formed through the spontaneous inter-specific hybridization of turnip-rape (*Brassica rapa*, 2n= 20) and cabbage (*Brassica oleracea*, 2n=18) (Chalhoub *et al*. 2014). The genetic diversity among the modern breeding material of canola is low (Gomez 1999) which is mainly attributed to intensive breeding for specific traits such as oil and seed quality (Hasan *et al*. 2006). More recently canola breeding efforts have shifted to the development of hybrids. Hybrid breeding is a method of selecting parental lines and determining crosses that are likely to outperform their inbred progenitors. Although hybrid breeding requires more time and resources compared to inbred breeding (Cros et al, 2018), it is preferred as the hybrid varieties can outperform their inbred progenitors (Fu et al 2014).

In canola male and female heterotic groups follow different pipelines, where the female inbred line development goes through a male sterility procedure. Cytoplasmic male sterility (CMS) in the female parental line is achieved through Ogura CMS (ogu CMS)(Ogura, 1967). This process adds one to two extra years to the development of female lines compared to the male inbred lines. For hybrid production, male lines carrying restorer nuclear genes are used to restore fertility in the hybrid plants. Due to the resource intensive nature of effective introgression of Ogura CMS, heavy selection is often placed on the female heterotic pool prior to CMS conversion based on limited inbred performance data. Genomic prediction offers a promising approach to improve the accuracy and efficiency of selecting female lines to undergo CMS conversion, without the need for inbred testing.

Genomic prediction is a tool that utilizes high density genetic markers that are in linkage disequilibrium with traits of interest to predict the genetic values of untested candidates based on their genotypic information (Meuwissen *et al*. 2001). Several studies have found that the use of genomic information can increase the accuracy of estimating additive genetic values (Crossa *et al*. 2014; Würschum *et al*. 2014; Zhao *et al*. 2015), and the use of genomic selection increases the rate of genetic gain when compared to classical phenotype-based selection methods (Zhao *et al*. 2015; Zou *et al*. 2016). While genomic prediction is a promising approach for improved breeding efficiency, prediction accuracy can be affected by the genetic architecture of the trait, heritability, the size of training set and the relatedness of training set with the validation set (De Los Campos *et al*. 2009; Terraillon *et al*. 2023).

In hybrid breeding, genomic prediction can be used for the prediction of general combining ability (GCA) as well as specific combining ability (SCA) of the parents. Genomic prediction models are trained using existing data to estimate the inbred parents combining abilities (Bernardo and Yu 2007) with GCA being additively inherited and SCA explaining cross specific deviations from the GCAs values of the parents. Hybrid breeding is designed for the exploitation of heterosis, which is the superior performance of F1 hybrids relative to the inbred parent performance (Schulthess *et al*. 2017). Development of genetically distant heterotic groups depends on the availability of high genetic diversity. In canola the genetic diversity in the modern breeding material is narrow (Basunanda *et al*. 2010) restricting the development of genetically distant heterotic groups. Given the relatively recent focus on hybrid breeding in canola, when compared to other crops such as maize, improving the efficiency of selecting parents in canola for better performing hybrids is crucial for the development of high performing hybrid lines. Studies (Jan *et al*. 2016; Zou *et al*. 2016) have shown the promising potential of genomic prediction in canola, but prediction models and genomic selection approaches need to be optimized for the canola hybrid breeding system.

The goal of the present study was to evaluate population structure, determine genetic correlations between inbred per se and hybrid GCA for the female heterotic pool, estimate (co)variance parameters for commercially important traits, and evaluate the accuracy of genomic prediction in a commercial canola breeding program. Single step Genomic best linear unbiased prediction (GBLUP) models were implemented, and prediction accuracy was assessed in three different cross validation strategies. In the first cross validation (CV1) strategy, five-fold cross validation was applied while in the second cross-validation (CV2) strategy, prediction was done for the unseen year. In the third cross-validation (CV3) strategy, inbred per se performance was added to the training set to exploit the genetic co-variance between hybrid progenitors and hybrid data.

## Material and Methods

### Field Trial

The phenotypic data used in this study were collected from commercial field trials conducted by Nutrien Ag Solutions over four years (2018 - 2021) and across 12 locations in Canada. Trials were defined as groups of genotypes tested together in multiple experimental designs. In total, there were 47 line by tester trials for testing female hybrid parent performance and 28 line by tester trials for testing male hybrid parent performance. For the female line by tester program, hundreds of female lines are testcrossed with selected male lines while for the male line by tester program hundreds of male lines are testcrossed with selected male tester lines. The trial design was a combination of unreplicated and partial replicated trials, where trials are connected by checks and genetic relationships. Details of the female and male line by tester trials are described in supplementary table 1.

### Phenotype data

Phenotypic data was collected for seven traits including grain yield, seed quality, and morphological traits. Grain yield (GY) in kg per hectare was calculated by adjusting the total seed weight per plot to 8.5% moisture and converted to kg per hectare. Seed oil content (%) and protein content (%) were measured as the percentage of oil and protein content in the total seed dry matter. Total glucosinolate (TGLU) content within the seed was expressed as micromoles per gram (µmoles/g) of whole seed and adjusted to 8.5% moisture. Total saturated fatty acids (SAT) were measured as a percentage of total fatty acids in a seed. Oil, protein, total glucosinolate (TGLU) and saturated fat (SAT) were measured using a FOSS near-infrared reflectance spectroscopy System 6500 instrument (FOSS, Hillerod, Denmark, pre-calibrated with the analysis data of the same samples using gas chromatography). Days to flowering (DTF) was recorded as the number of days that have elapsed between the planting date and the date when 50% of the plants in the plot have at least one flower open and days to maturity (DTM) were recorded as the number of days that have elapsed between the planting date and the date when seed coat color change is observed halfway up the main raceme in 50% of the plants in a plot. The phenotype data was cleaned by removing data records outside ±3 standardized residuals and lines with no genomic information were removed from the analysis. Descriptive statistics of the traits are presented in supplementary table 2.

### Genotyping

Genotyping was done using Diversity Array Technology Pty Ltd (DArT, University of Canberra, Bruce, ACT Australia) genotyping platform. DNA extraction was done on 10 seed samples per female double haploid (DH) parental line. DNA extraction was done following DArT protocols (https://www.diversityarrays.com/services/laboratory-services/). Initial genotyping was carried out using genotyping-by-sequencing-based DArTseq methods for SNP discovery purposes, and a 2000 mid density SNP panel was used for genotyping hybrid parent lines. Quality control was performed by removing SNPs with a call rate lower than 0.90, heterozygote frequency higher than 0.10 and minimum allele frequency (MAF) lower than 0.05. Markers and individuals with more than 20% missing marker data were removed. Remaining missing genotype data was imputed using mean genotype dosage for each marker. In total, 1365 SNPs passed the quality control, and the cleaned genotype data was used to construct genomic relationship matrix *G* using the A.mat function from rrBLUP R package (Endelman 2011; VanRaden 2008). The parent genomic relationship matrix *G*_GCA_was computed with the following formula:

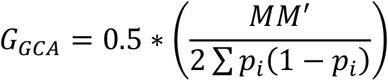

Where *M* is the centered genotypic matrix (genotypes were represented as counts of the reference allele: 0, 1, 2), and *p*_i_ is the minor allele frequency of *i*^th^ SNP. For checks where the parents were not genotyped, they were included in the GRM with a diagonal value of one 1 and off-diagonal values of zero.

### Projecting hybrid genotypes

Hybrid genotypes were derived from genotypes of the corresponding inbred lines. If both female and male parents are homozygous for the same allele, then the F1 progeny receives homozygous. If the parental line were homozygous for opposite alleles at a given marker, then the F1 progeny received a heterozygous genotype. For the parental genotype, the heterozygosity was kept less than 10%, and for these heterozygous parents the F1 progeny received the mean of the parents. The dominance relationship matrix *G*_D_ was computed using the following formula:

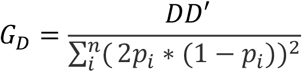

Where *D* is the centered genotypic matrix, with specific coding for homozygous and heterozygous individuals at each locus.

### Statistical models

Single stage genomic best linear unbiased prediction (GBLUP) model was implemented to estimate variances contributed by female and male parents as well as hybrid effect. The combined analysis was done to determine partitioning of genetic variance between GCA and SCA. For evaluation of genomic prediction accuracy, focus was placed on the female germplasm pool due to a lack of connectivity between the male and female tester datasets, and the importance of the female germplasm pool in the hybrid testing.

1. **GCA + SCA model**

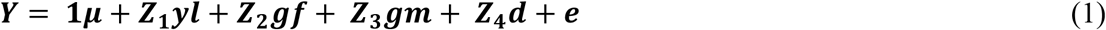

Where ***y***_***n***_ is the vector of observations (grain yield, oil, protein, days to flowering, days to maturity, TGLU, and SAT). μ is an intercept, **Z**_**1**_, **Z**_**2**_, **Z**_**3**_, and **Z**_**4**_ are the design matrices of random effects; ***yl*** is a vector of nested year, location and layout effects; ***gf*** is the vector of GCA effects for female, 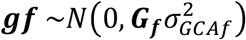, where ***G***_***f***_ is the genomic relationship matrix and 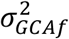 is the female GCA genomic variance; **g*m*** is the vector of GCA effects for male, 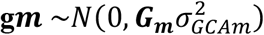, where ***G***_***m***_ is the genomic relationship matrix for male inbred parents and 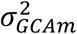 is the genomic variance for the male parent GCA effect; ***d*** is the vector of SCA effects for hybrid, 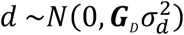, where ***G***_***D***_ is the dominance relationship matrix and 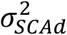 is the hybrid SCA genomic variance; *e* is the vector of random residual effect with 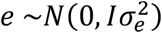, ***I*** is an identity matrix and 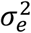 is the residual variance.
2. **Female GCA model** The following model was implemented for the prediction of female GCA for the first and second cross-validation strategy.

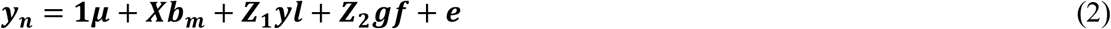

Where ***y***_***n***_ is the vector of observations, as described earlier. μ is an intercept, ***X*** is the design matrix for the fixed effects; ***b*** is the vector of fixed effect of male tester; **Z**_**1**_, ***yl, f, e*** are as described in the earlier model.
3. **Female inbred hybrid covariance model** This model was implemented in the third cross validation scenario (Figure 1) that exploits the genetic co-variance between inbred per se performance and inbred GCA.

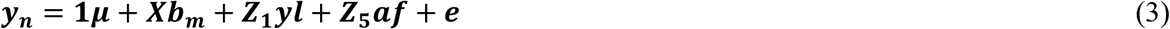 All the model terms are similar as the previous model, except for the variance structure of*a* and *e*. Where ***af*** is the vector of GCA effects for female inbred per se performance and GCA, ***af*** ∼ ***N***(**0, *G***_***f***_**⨂*G***_**0**_), 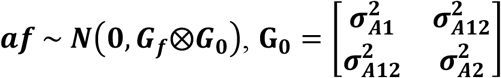 where ***G***_***f***_ is the additive relationship matrix and ***G***_0_ is unstructured variance-covariance matrix for inbred and hybrid effects and **⨂** is the Kronecker product. The diagonal elements of ***G***_O_ are the additive genetic variance of inbred per se and inbred GCA, while the off-diagonal elements are the genetic covariance between inbred per se and inbred GCA performance. The error was distributed as 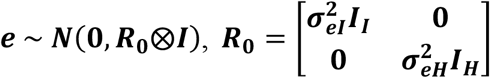. Where ***R***_**0**_ is the residual variance-covariance and 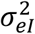 and 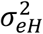 are error variance of inbred per se and hybrid phenotypes. *I*_l_ and *I*_H_ are identity matrices and residuals are assumed to be uncorrelated between inbred and hybrid trials.

**Figure 1.**
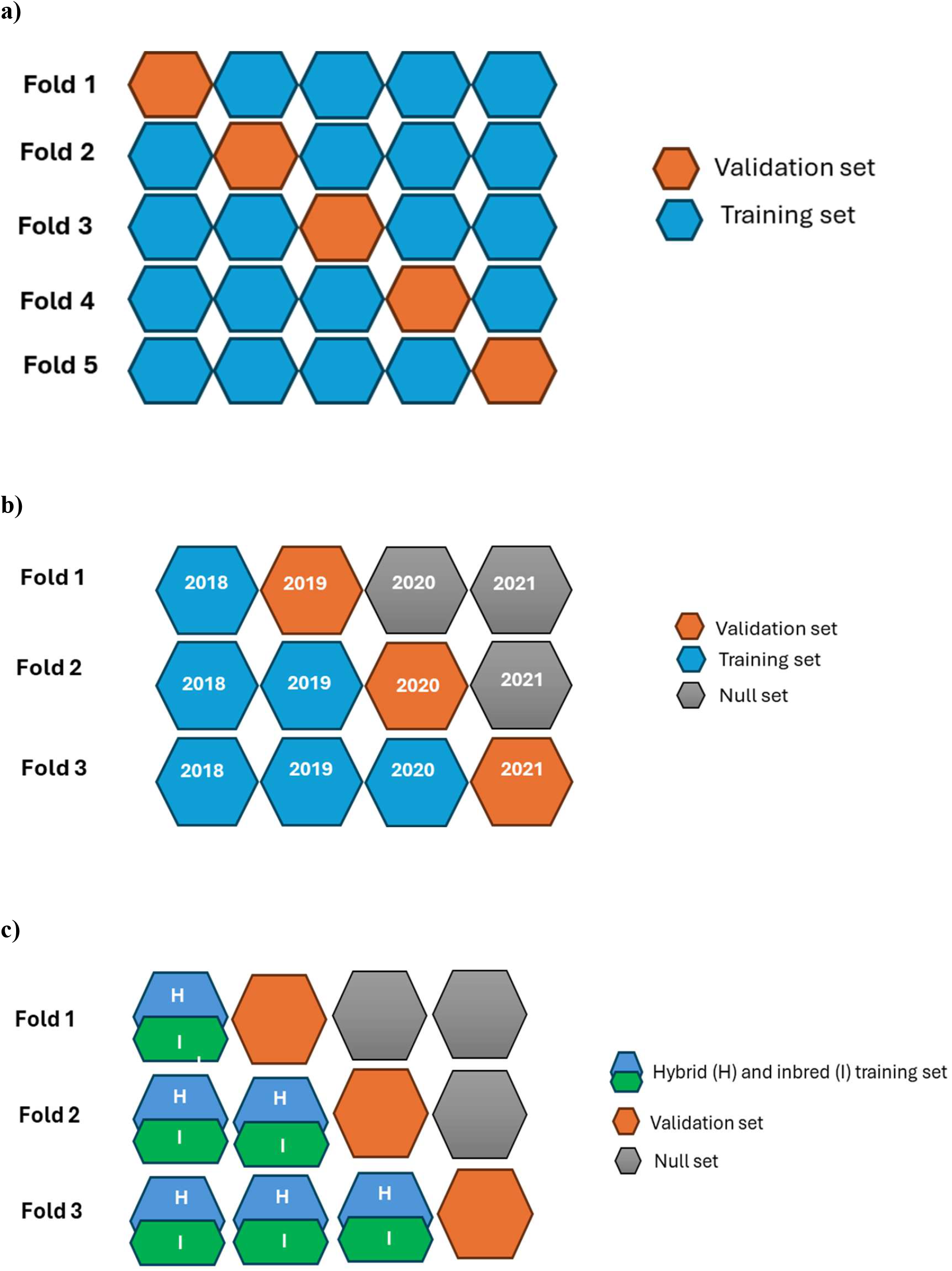
schematic representation of cross-validation strategies; a) five-fold cross cv (CV1), b) untested-line cv (CV2) and c) inbred-hybrid covariance cv (CV3). Null set in CV2 and CV3 is data set that was not included in the training set.

### BLUE estimation

A best linear unbiased estimate (BLUE) was made for each trait as:

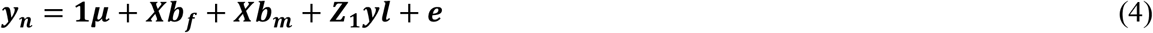

Where ***y***_***n***_ is the vector of observations of traits listed previously. μ is an intercept, ***X*** is the design matrix for the fixed effects; ***b***___ are the vectors of fixed effects for female *b* and male *b*_m_; where **Z**_**1**_, ***yl, e*** are same as described earlier. The BLUEs were used for cross validation by correlating predicted performance with BLUEs estimated form the complete dataset.

### Variance components and heritability

The restricted maximum likelihood (REML) method was used to estimate variance components for calculation of heritability for traits collected in hybrid trials using ASREML command line function for version 4.1(Gilmour et al., 2015). The following model was implemented to estimate variance components for broad sense heritability.

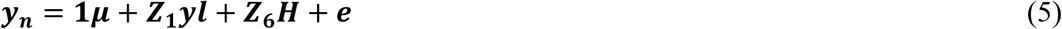

Where ***y***_***n***_ is the vector of observations of traits listed previously, μ is an intercept, **Z**_6_ is the design matrix for the random effect of hybrid; ***yl*** is a vector of year, location and layout nested within each other; ***e*** is the vector of random residual effect with 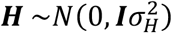, where ***I*** is an identity matrix and 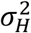 is the hybrid variance.

Broad sense heritability 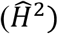 was estimated with the following formula:

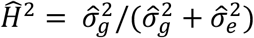

where 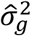 is the genetic variance, 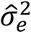 is the estimated residual variance.

### Cross-validation schemes and model validation

We tested three different cross-validation schemes, to examine the performance of genomic prediction models using historical inbred and hybrid canola testing data. In the first cross-validation scheme (CV1), the prediction model was validated using a five-fold cross validation strategy. Phenotypic data was split into five sets, where sets were designed to ensure all hybrids from a given set of parents were included in the same set. This was done to avoid inflating prediction accuracy by having hybrids with the same parent(s) in both the training and test sets. The models were trained on four folds to predict a fifth fold that was removed from the training set. This process was repeated until all sets were predicted.

For the second cross-validation scheme (CV2), prediction was done for the GCA of female hybrid parents, and this was achieved by exploiting the genetic covariance between female hybrid genetic covariance between tested and untested hybrids. The genomic relationship matrix was used to estimate the covariance between tested and untested female hybrid parents. Four years (2018 – 2021) of data was available and genomic prediction was carried out for the three years (2019 – 2021). For predicting 2019, the training set was formed using 2018 data and for predicting 2020, hybrid data from 2018 and 2019 was used for the training set. For the prediction of 2021, hybrid data from 2018 to 2020 was used in the training set.

The third cross-validation scheme (CV3) exploits the covariance between traits of inbred per se and parental GCA performance. The covariance was estimated using the SNP markers and observed inbred per se and hybrid performance data. Inbred per se performance data was available from 2015 to 2019 for building a training set. Given inbred per se performance data is collected prior to CMS conversion, data collected 2-years prior to corresponding hybrid performance trials was included to determine the impact of including inbred per se performance data on the accuracy of genomic predictions for inbred parent GCA. For predicting 2019, inbred data from 2015 to 2017 and hybrid data of 2018 was used to train the model. For predicting 2020, inbred data from 2015 to 2017 and hybrid data of 2018 and 2019 was used to build the training set. For the prediction of 2021, inbred data from 2015 to 2019 and hybrid data from 2018 to 2020 was used to train the model.

Predictive ability (PA) of the models was computed as the Pearson correlation between BLUE values (ŷ _BLUE_) and the predicted values (ŷ _prediction_).

## Results

### Population structure

Principal component analysis of the female and male heterotic group is presented in Figure 2. The first PC explained 8.21% while PC 2 explained 5.91% of genetic variation. Plots of the first two principal components show that, while PC1 does differentiate male and femal heterotic groups, there is not clear separation. This was in line with previous studies that heterotic groups in canola are not genetically diverse (Jan *et al*. 2016).

**Figure 2.**
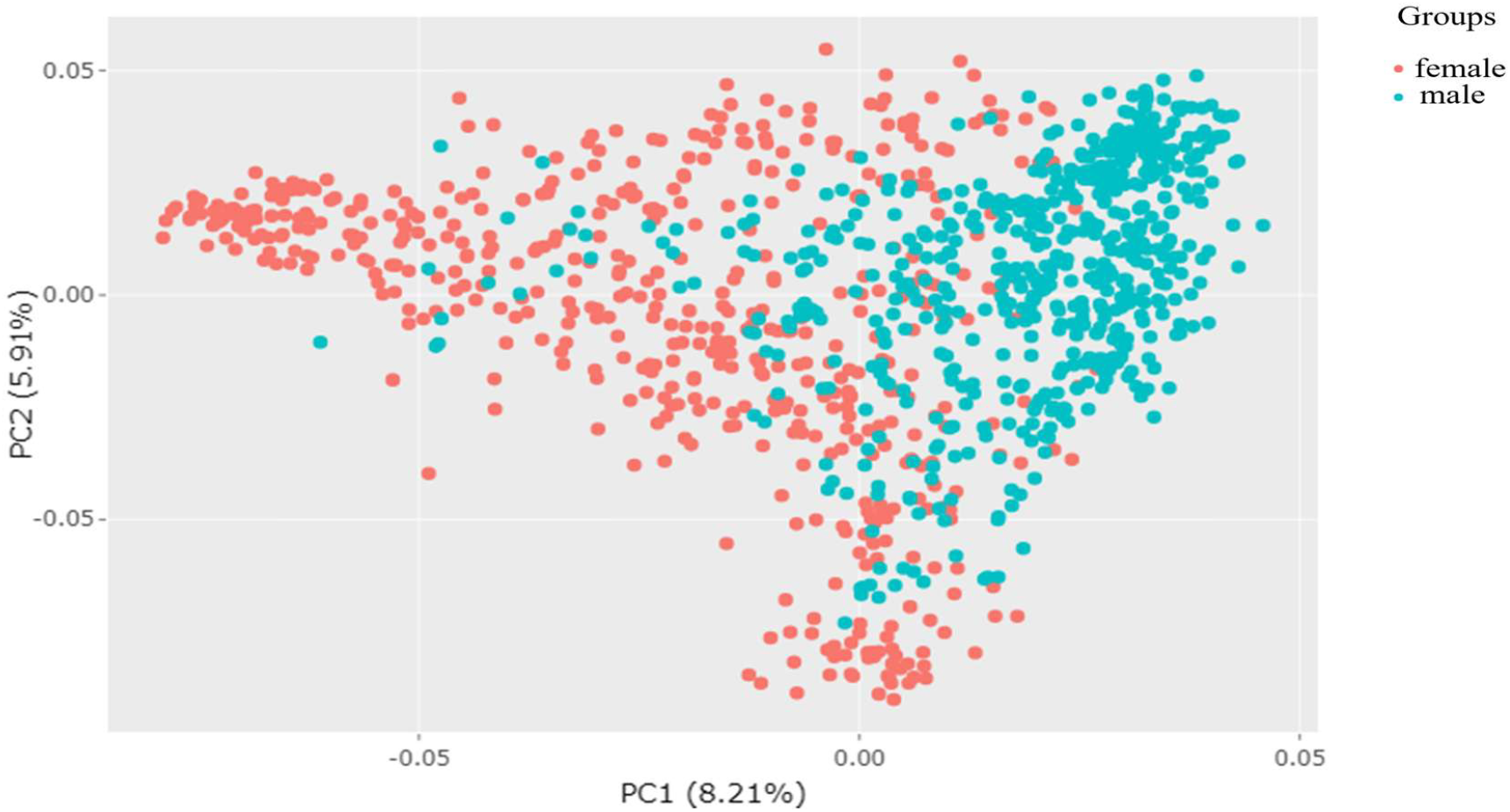
Principal component analysis (PCA) of male and female heterotic groups of canola using 1365 SNPs.

**Figure 3.**
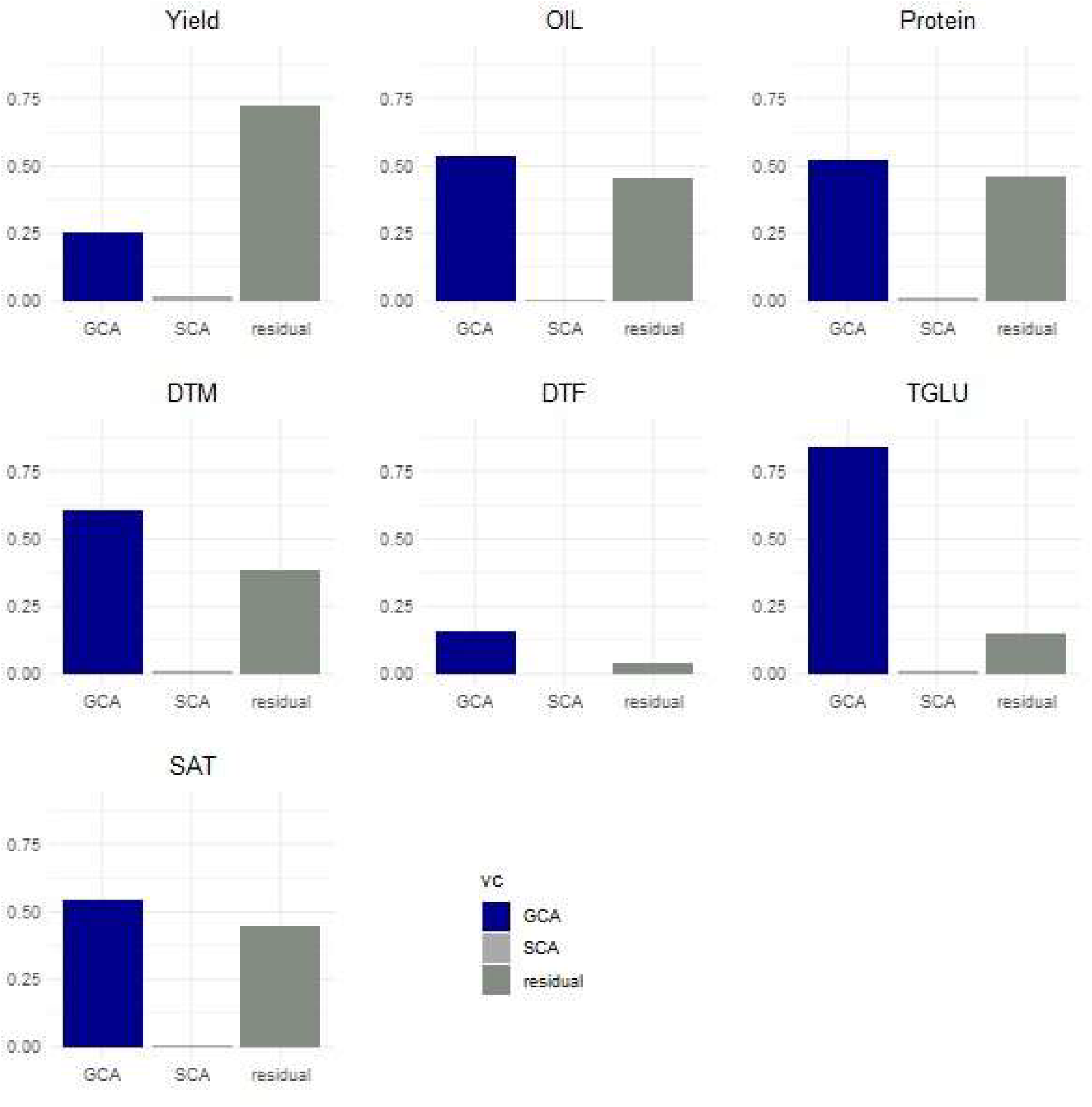
proportion of variance component from GCA + SCA model. GCA: general combining ability, SCA: specific combining ability.

### GCA and SCA model

The proportion of genetic variance explained by GCA and SCA is presented in Figure 4. For all the traits studied, genetic variance was explained by primarily by GCA, with SCA explaining a small portion of the total genetic variance. The highest proportion of total variation explained by GCA (83.8%) was observed for TGLU while the lowest proportion of variation explained by GCA (15.9%) was observed for DTF. The genetic proportion explained by GCA for Protein and oil was 52.6% and 53.9%, respectively. GCA explained 60.3% for DTM, 54.7% for SAT, 25.4% for yield. The proportion of non-genetic variance, which is the residual variance was higher for yield. The genetic variance explained by SCA was less than 1% for all traits. Further investigation with the GCA and SCA model was not performed given the negligible proportion of variance explained by SCA.

**Figure 4.**
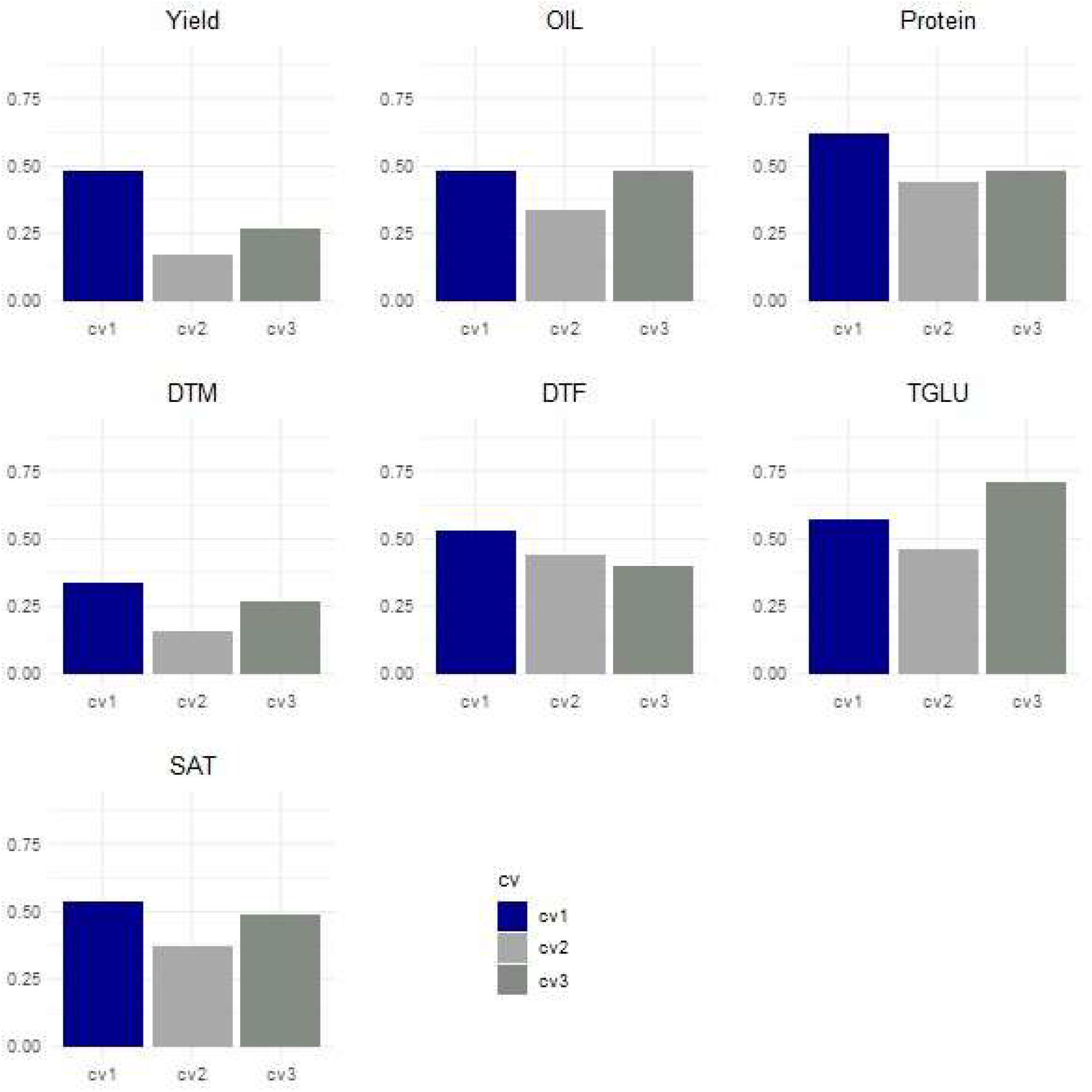
Bar plot of predictive ability for the three different cross-validation strategies for the seven traits. CV1 = five-fold cross validation, CV2 = untested line prediction, CV3 = untested line prediction with inbred hybrid covariance. DTM is days to maturity, DTF is days to flowering and SAT is saturated fatty acid.

### Variance components, broad sense heritability and genetic correlation

The variance components, broad sense heritability along with the summary statistics is presented in Table 1. The genetic 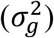 and residual variances 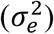 were estimated for the hybrid effect (Eq.5). This was done to capture the genetic variance contributed both by female and male parents in the female tester data, where hundreds of females are crossed with about six male testers. Broad sense heritability ranged 0.15 to 0.53 across the measured traits. Higher *H*^2^ was estimated for TGLU and DTF (0.53) followed by SAT (0.37), while lower *H*^2^ estimate was reported for grain yield (0.15). The genetic correlation between inbred and hybrid female tester data ranged from 0.45 to 0.88. The highest genetic correlation was for TGLU (*r* = 0.88) while the lowest genetic correlation was for oil (*r* = 0.45).

**Table 1.** Summary statistics of seven different traits of female tester data used for genomic prediction analysis.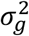: genetic variance, 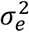: residual variance, *H*^2^ : Broad sense heritability, *r*: genetic correlation of traits collected under inbred trial and hybrid trial.

| Traits | $\sigma_g^2$ | $\sigma_e^2$ | $H^2$ | $r$ |
| --- | --- | --- | --- | --- |
| Grain yield (kg/ha) | 23428.6<br>(1452.48) | 126473 (1757.05) | 0.15 | 0.69 |
| Oil (%) | 0.82 (0.03) | 1.61 (0.02) | 0.33 | 0.45 |
| Protein (%) | 1.13 (0.05) | 2.33 (0.03) | 0.32 | 0.73 |
| Days to flowering (DAP) | 1.48 (0.06) | 1.27 (0.03) | 0.53 | 0.8 |
| Days to maturity (DAP) | 1.48 (0.06) | 3.85 (0.05) | 0.27 | 0.84 |
| TGLU (μmoles /g) | 2.03 (0.07) | 1.77 (0.02) | 0.53 | 0.88 |
| SAT (%) | 0.017 (0.0007) | 0.028 (0.0004) | 0.37 | 0.62 |

### Genomic prediction

The predictive ability for the three different cross validation strategies studied is presented in Figure 1. The three different cross-validation strategies investigated were, CV1-five-fold cross validation, CV2-prediction of untested lines and CV3-cross validation of untested lines with inbred-hybrid covariance. The highest PA was observed for CV1 while the lowest PA was seen for CV2. In the CV1, PA ranged from 0.34 to 0.62. The highest PA was observed for protein (0.62) while the lowest PA was observed for DTM (0.34). For CV2, PA ranged from 0.16 to 0.46, while for CV3 PA ranged from 0.27 to 0.71. The highest PA for CV2 (0.46) and CV3 (0.71) was observed for TGLU while the lowest PA (0.16 – CV2 and 0.27 – CV3) was for DTM.

## Discussion

In the current study, three different cross-validation strategies were evaluated to estimate general combining ability of potential female parents within a commercial canola hybrid breeding program. The predictive ability for each cross-validation strategy showed different values as shown in Figure 4. CV1 has highest PA for most of the traits compared to CV2 and CV3. CV1 was validated using five-fold which would have more relatedness between training and validation sets compared to CV2 and CV3. CV2 and CV3 predicted untested lines with the only difference between CV2 and CV3 being that inbred data was added in the training set for CV3. For all the traits, CV3 has a better predictive ability than CV2 and this can be explained by the high genetic correlation of traits between inbred and hybrid which ranged from 0.45 to 0.88.

### Variance components, heritability, and genetic correlation

In the GCA and SCA model, a significant proportion of the genetic variance was captured by the GCA of female and male parents while variance explained by SCA was very low. The additive genetic variance was predominant compared to the non-additive variance. This could be observed in the PCA plot, where the male and female heterotic groups did not show clear distinction (Figure 2). This result was also in line with other studies (Jan *et al*. 2016) that indicated heterotic groups in canola are not genetically distant. This indicates there is a need to select for divergent founders when developing heterotic groups to help better exploit heterosis. Divergent founders increase the ratio of GCA to SCA and dominance is expected to be positively correlated with increasing genetic distance of parents (Reif *et al*. 2007). The genetic correlation between traits measured under inbred and hybrid female tester trials showed medium to high correlation. The genetic correlation helps to exploit the genetic covariance between inbred and hybrid trials, and this was shown in our prediction result where the training set with inbred data outperformed those without inbred data.

### Genomic predictive ability

In the current study, sufficient PA for most of the traits indicates the potential of genomic selection in canola hybrid breeding. The PA of GCA for grain yield in the current study was 0.48 for CV1, 0.17 for CV2 and 0.27 for CV3. The higher PA for CV1 can be explained by the higher relatedness of training and validation set in CV1 compared to CV2 and CV3. Relatedness between training and validation set is one of the factors that affects prediction accuracy (De Los Campos *et al*. 2009) and high relatedness between training and validation sets has been reported to increase prediction accuracy (Habier *et al*. 2007; Hayes *et al*. 2009). In addition to the difference in relatedness, the training size between CV1 and CV2 is different (Figure 1). In CV2 data from the previous year are available to train the model which resulted in a smaller training set compared to CV1. However, the training set size increased by 168.65 % from the first-year prediction when predicting the last year. A prediction accuracy of 0.45 for yield in canola test cross was reported (Jan *et al*. 2016)in a study that investigated genomic prediction of hybrid performance. However, the current study was focused on GCA of the hybrid parents. Information on the GCA of parents plays a significant role in a breeder’s decision-making process to select female parents for CMS conversion and male parents for superior hybrid production. Besides seed yield, quality traits are another important trait in canola breeding (Jan *et al*. 2016; Zou *et al*. 2016). The PA of these quality traits ranged from 0.48 to 0.62 for CV1. Seed quality traits represented in the current study were seed oil content (PA= 0.48), SAT (PA = 0.54), TGLU (PA = 0.57), and protein content (PA = 0.62). Prediction accuracies of 0.41 for glucosinolate content and 0.34 for yield have been reported in canola (Würschum et al. 2014). The PA for DTF and DTM were 0.53 and 0.34, respectively. (Würschum *et al*. 2014) also reported prediction accuracy of 0.8 for flowering time, and 0.6 to 0.7 for protein and oil content.

When predicting untested lines, the PA of CV3 outperformed CV2 for all the traits. In CV3, the training set was extended with the addition of inbred per se performance data which could explain why CV3 has improved predictive ability compared to CV2. The genetic correlation between inbred and hybrid trials was observed to be high for all the traits (Table 1). The covariance between inbred and hybrid trials contributed to the improvement of PA and this was in line with the study of (Liang *et al*. 2018) in pearl millet. However, in their study correcting for heterosis per se gave a better prediction accuracy than incorporating inbred phenotypic data without correcting for heterosis. In the current study, the contribution of SCA variance is very low. The incorporation of inbred data has showed an increment in PA that ranged from 9 – 68.7%. The highest increase was for DTM followed by GY, while the least increase was observed for protein content.

### Implementing genomic prediction in canola breeding

Unlike crops such as maize, which exhibit significant genetic divergence between heterotic pools, canola’s heterotic pools are relatively less genetically distinct (Jan et al. 2016). This was also demonstrated in the current study (Fig. 2). This narrow genetic base presents both challenges and opportunities for genomic breeding in canola. The lack of genetic diversity between heterotic pools may limit the chance of exploiting heterosis. On the other hand, the lack of genetic diversity provides an opportunity to implement genomic prediction strategies that could enhance the accuracy of parent selection and improve hybrid performance. The integration of molecular markers into breeding programs has revolutionized the selection process, offering a more accurate and faster alternative to traditional phenotype-based selection. In canola, studies by (Jan et al. 2016; Zou et al. 2016) have demonstrated the potential of genomic prediction for improved selection accuracy. Accurate prediction of the GCA of parents can contribute to the improvement of genetic gain in hybrid canola breeding. Other studies (Crossa et al. 2014; Würschum et al. 2014; Zhao et al. 2015) have shown the advantages of using molecular markers over classical phenotype selection in improving selection accuracy. Besides improving selection accuracy, the integration of genomic breeding strategies in canola also provides an opportunity to shorten the breeding cycle.

In conclusion, genomic prediction represents a promising strategy for improving hybrid canola breeding. The ability to accurately estimate GCA of parents allows breeders to make better informed decisions. Our results have also shown the potential of genomic prediction to enhance breeding efficiency in canola breeding.

## Data availability

The dataset used in the current study are available in the repository https://github.com/Biruk2B/canola_paper

We would like to thank Nutrien Ag Solutions for providing the genotype and phenotype data as well as funding this work.

## Author Contributions

BBT: conceptualization, data curation, methodology, analysis, writing. CPA, KP, KY, ZE, TK, TA, KL, WB: resources, phenotyping and genotyping. KR: conceptualization, funding acquisition and supervision. All authors have read and reviewed the manuscripts.

