## supplementary table 1 for "Genomic prediction for general combining ability in hybrid canola (*Brassica napa* L.)"

### Supplementary material

**Supplementary table 1 Female line by tester phenotype data collected from 2018 to 2021 across 12 different locations. Number of plots, female lines, male testers and hybrids created are shown.**

| Year | # plots | Female | Male | # unique hybrids |
|---|---|---|---|---|
| 2018 | 3700 | 127 | 6 | 619 |
| 2019 | 1298 | 74 | 5 | 264 |
| 2020 | 4936 | 133 | 6 | 664 |
| 2021 | 3899 | 116 | 6 | 565 |
